# GI-16 lineage (624/I or Q1), there and Back Again: the history of one of the major threat for poultry farming of our era

**DOI:** 10.1101/402800

**Authors:** Giovanni Franzo, Mattia Cecchinato, Giovanni Tosi, Laura Fiorentini, Francesca Faccin, Claudia Maria Tucciarone, Tiziana Trogu, Ilaria Barbieri, Paola Massi, Ana Moreno

**Author notes:** Corresponding authors:* Giovanni Franzo, Department of Animal Medicine, Production and Health (MAPS), University of Padua, Viale dell’Università 16, 35020 Legnaro (PD), Italy.; Ana Moreno, Department of Virology, Istituto Zooprofilattico Sperimentale della Lombardia ed Emilia Romagna, Via A. Bianchi, 9 25124, Brescia (BS) Italy.

## Abstract

The genetic variability of Infectious bronchitis virus (IBV) is one of the main challenges for its control, hindering not only the development of effective vaccination strategies but also its classification and, consequently, epidemiology understanding. The 624/I and Q1 genotypes, now recognized to be part of the GI-16 lineage, represent an excellent example of the practical consequences of IBV molecular epidemiology limited knowledge. In fact, being their common origin unrecognized for a long time, independent epidemiological pictures were drawn for the two genotypes. To fix this misinterpretation, the present study reconstructs the history, population dynamics and spreading patterns of GI-16 lineage as a whole using a phylodynamic approach. A collection of worldwide available hypervariable region 1 and 2 (HVR12) and 3 (HVR3) sequences of the S1 protein was analysed together with 258 HVR3 sequences obtained from samples collected in Italy (the country where this genotype was initially identified) since 1963. The results demonstrate that after its emergence at the beginning of the XX century, GI-16 was able to persist until present days in Italy. Approximately in the late 1980s, it migrated to Asia, which became the main nucleus for further spreading to Middle East, Europe and especially South America, likely through multiple introduction events. A remarkable among-country diffusion was also demonstrated in Asia and South America. Interestingly, although most of the recent Italian GI-16 strains originated from ancestral viruses detected in the same country, a couple were closely related to Chinese ones, supporting a backward viral flow from China to Italy.

Besides to the specific case-study results, this work highlights the misconceptions that originate from the lack of a unified nomenclature and poor molecular epidemiology data generation and sharing. This shortcoming appears particularly relevant since the described scenario could likely be shared by many other IBV genotypes and pathogens in general.

## Introduction

Infectious bronchitis virus (IBV) is currently classified in the species *Avian coronavirus*, genus *Gammacoronavirus*, family *Coronaviridae.* The viral genome is about 27 Kb long and encodes for different proteins, such as the RNA-dependent RNA polymerase (RdRp), numerous accessory and regulatory proteins and the Spike, Envelope, Membrane, and Nucleocapsid structural proteins [1]. IBV currently represents one of the most relevant diseases of poultry farming, being responsible for respiratory signs, reproductive disorders and relevant mortality, particularly in presence of nephropathogenic strains or secondary infections [2]. Similarly to other RNA viruses, it is characterized by high evolutionary and recombination rates, which drive the rapid evolution of this pathogen and lead to a remarkable genetic and phenotypic heterogeneity [3]. The Spike protein, further processed in the S1 and S2 sub-units, is by far the most studied viral protein because of its relevance in viral life-cycle and host interaction. Besides being involved in viral attachment and viral-host membranes fusion, it is also the main target of the host neutralizing immune response [1,4]. Its remarkable variability has several implications, the most practical one being the poor cross protection among different viral strains and genotypes [5], which severely hinders the infection and disease control through vaccination. Additionally, the genetic variability has also complicated the viral classification, leading to the proposal of a multitude of genotypes based on specific research studies or local investigations, in absence of shared criteria. Although it could appear a mere nomenclature issue, this state of affairs had astonishing implications impeding or confusing the proper understanding of IBV molecular epidemiology.

Only in 2016, in the framework of COST Action FA1207, a coherent and common IBV genotype classification was proposed [6], grouping the identified strains in 6 genotypes and 32 lineages. Interestingly, Valastro et al., (2016), confirmed what previously proposed by Franzo et al., [7] about the clustering of 624/I and Q1 genotypes in the same group (currently named GI-16). Until then, the two genotypes had been named (and considered) independent: the 624/I was first reported in Italy in 1993 [8], and thereafter in Slovenia [9], Poland [10], and Russia [11]. Moreover, retrospective studies demonstrated its wide circulation in Italy since the early 1960s [12], where it remained the most prevalent genotype for more than a decade. However, from the 1990s, its detection and reporting became sporadic or absent. At the same time, in 1996, a new virus (later renamed Q1) was identified in animals affected by proventriculitis [13] in China and thereafter in other Asian [14,15], Middle Eastern [16,17], South American [18,19] and European countries [7,20–22]. Remarkably, not recognizing the relationship between these strains, a new spreading from Asia was supposed, similarly to what proven for the GI-19 lineage (QX genotype) [23]. The insight provided by recent genetic studies [6,7] raised new questions about the history, epidemiology and spreading of the GI-16 lineage around the world. The knowledge of these processes would be of interest not only for the particular case under investigation, but could also provide useful information on the general intra- and inter-continents viral dispersion patterns, aiding the development and implementation of adequate control measures.

Unfortunately, the paucity of sequences available before Q1 detection in China prevented this investigation for a long time. In the present study, a high number of retrospective GI-16 sequences were generated from Italian samples collected since 1963 and analyzed together with the worldwide available ones using a phylodynamic approach to reconstruct the history of this genotype and infer its spreading pattern around the world.

## Material and Methods

### Italian strain detection and sequencing

IBV samples collected between 1963-2017 were included in the study. Italian sequences were obtained during the routine diagnostic activity of the Istituto Zooprofilattico Sperimentale della Lombardia e dell’Emilia Romagna. Samples of trachea, kidneys, lungs, cecal tonsils, or tracheal swabs were delivered by field veterinarians from the whole country but most of them originated from the highly densely populated regions of Northern Italy. Of these, samples obtained between 1963-2009 were previously isolated in chicken SPF eggs and were available in the archives of Istituto Zooprofilattico Sperimentale della Lombardia e dell’Emilia-Romagna. The sequencing of samples collected before 2009 was performed on the isolates whereas the sequencing of those collected after 2009 was directly carried outon the submitted tissues..

Viral RNA was extracted from isolates using the TRIzol reagent (Invitrogen, Carlsbad, CA, USA) according to the manufacturer’s protocol. IBV was detected using RT-PCR as previously described [23]. Partial sequencing of the S1 gene (464-bp), covering the hypervariable region 3 (HVR3), was performed using the same primers that were used for RT-PCR. The obtained chromatogram quality was evaluated by FinchTV (**http://www.geospiza.com**) and the consensus sequences were reconstructed using CromasPro (CromasPro Version 1.5).

### International database preparation

To download all available IBV sequences overlapping with the region of interest (HVR3), one of the Italian sequences was used as query in BLAST and all sequences with a genetic distance lower than 30% were downloaded. Obtained sequences plus the Italian ones were merged with the reference set provided by Valastro et al., [6] and aligned using MAFFT [24]. Partial sequences (i.e. covering only part of the HVR3) and those with unknown nucleotides, frameshift mutations and premature stop codons, suggestive of poor sequence quality, were removed from the dataset. A phylogenetic tree was then reconstructed using PhyML [25] selecting as the most appropriate substitution model the one with the lowest Akaike Information Criterion (AIC) calculated using Jmodeltest [26]. The phylogenetic tree was used to classify the strains and those potentially belonging to the GI-16 lineage were extracted. The presence of recombinant strains was evaluated using RDP4 [27] on a dataset including the selected GI-16 like strains plus the reference ones. The RDP4 settings for each method were adjusted accounting for the dataset features according to the RDP manual recommendations. Only recombination events detected by more than 2 methods with a significance value lower than 10^-5^ (p-value < 10^-5^) and Bonferroni correction were accepted. After recombinant removal, the absence of residual breakpoints was confirmed with SBP implemented in HyPhy [28]. Only sequences with collection date and country were included in the study. Dedicated Python scripts were used to extract this information from the Genbank file or, if absent, original publications were consulted and the sequences were re-named accordingly. Finally, TempEst [29] was used to investigate the ‘temporal signal’ of the heterochronous sequences under investigation and the sequences whose sampling date was inconsistent with their genetic divergence and phylogenetic position were removed. Indeed, significant deviations from the expectation can be due to several factors including low sequencing quality, excessive passaging leading to the accumulation of cell-line adaptations, recombination, etc. [29]. Xia test was performed for each codon position to exclude the presence of codon saturation.

### Phylodynamic analysis

Since there was a clear bias in the number of sequences collected in each county and in different time periods, the original dataset was split in ten randomly generated databases in order to diminish this bias impact and evaluate the robustness of achieved results over several independent runs [30]. More specifically, a maximum of 5 sequences for each country-year pair were randomly selected and included in the study. For each dataset the time to Most Recent Common Ancestor (tMRCA), the evolutionary rates (substitutions/sites/year) and population dynamics (expressed as Effective population size (Ne)·generation time (t), also called relative genetic diversity) were jointly estimated in a Bayesian fashion using the serial coalescent approach implemented in BEAST 1.8.4 [31]. Additionally, the discrete state phylogeographic approach implemented in the same program was used to reconstruct the viral spreading pattern, assuming each collection country as a strain feature [32]. This method allows to estimate the location and time of ancestral strains, calibrated using the implemented molecular clock and population model. Above all, the Bayesian approach allows to account for all the other parameter and phylogenetic uncertainness. Additionally, the use of the Bayesian Stochastic Search Variable Selection (BSSVS) allows for a Bayesian Factor (BF) test that identifies the most parsimonious description of the spreading process [32]. The nucleotide substitution model (i.e. GTR+Γ_4_) was selected based on the results of Bayesian Information Criterion (BIC) calculated using Jmodeltest [26], while molecular clock (relaxed lognormal molecular clock [33]), population model (Bayesian skyline [34]) and phylogeographic substitution model (symmetric substitution model) were selected based on the BF value, calculated using the marginal likelihood of different models, estimated with the Path Sampling (PS) and Stepping Stone (SS) method [35].

For each dataset, a 200 million Markov Chain Monte Carlo (MCMC) chain was performed, sampling parameters and trees every 200.000 iterations. Run results, after discharging the first 20% of interactions as burn-in, were evaluated using Tracer 1.5 and accepted only if Estimated Sample Size (ESS) of each parameter was higher than 200 and run mixing and convergence were adequate. Parameter estimation were summarized and reported in term of mean and 95% Highest Posterior Density (HPD). Maximum clade credibility (MCC) trees were constructed and annotated using Treeannotator. SpreaD3 [36] was used to summarize and display phylogeographic results and to calculate the BF of different between-countries migration rates. The transmission rate was considered statistically significant (i.e. non-zero) when the BF was higher than 3 [32].

### Hypervariable region 1-2 (HVR12)

Unfortunately, no strict guidelines currently exist on IBV sequencing protocols. Consequently, despite the HVR3 of the S1 gene is the most commonly sequenced region, different research groups have produced and published sequences obtained from other Spike protein portions. To benefit from these sequences, some representative of additional countries, a second dataset based on the HVR1 and 2 (HVR12) was created and analyzed using the same approach previously described. However, because of the limited sequence number and more balanced origin, a unique analysis was performed, avoiding the multiple dataset generation step.

## Results

### Sequence dataset

A total of 258 Italian GI-16 HVR3 sequences were obtained, encompassing the period from 1963 to 2017 (Acc. Numbers MH705361 - MH705613). Globally, the final HVR3 dataset included sequences 310 sequences collected from 8 countries (S1 Table). TempEst analysis demonstrated the presence of a significant temporal structure, being the correlation between genetic distances and sampling dates equal to 0.54 (R^2^ = 0.29; slope 7.77×10^-4^).

The final HVR12 dataset included sequences 76 sequences collected from 12 countries (S1 Table). TempEst analysis demonstrated the presence of a significant temporal structure, being the correlation between genetic distances and sampling dates equal to 0.52 (R^2^ = 0.27; slope 2.70×10^-4^).

### Phylodynamic analysis HVR3

tMRCA of GI-16 genotype, averaged over all runs, was estimated in 1905.23 (1860.76-1934.36 95HPD), and the results were proven highly consistent among the different tested datasets (Figure 1). Similarly, repeatable results were obtained for the substitution rate analysis (Figure 1), which was estimated to be 1.89×10^-3^ (1.42×10^-3^–2.45×10^-3^).

**Figure 1.**
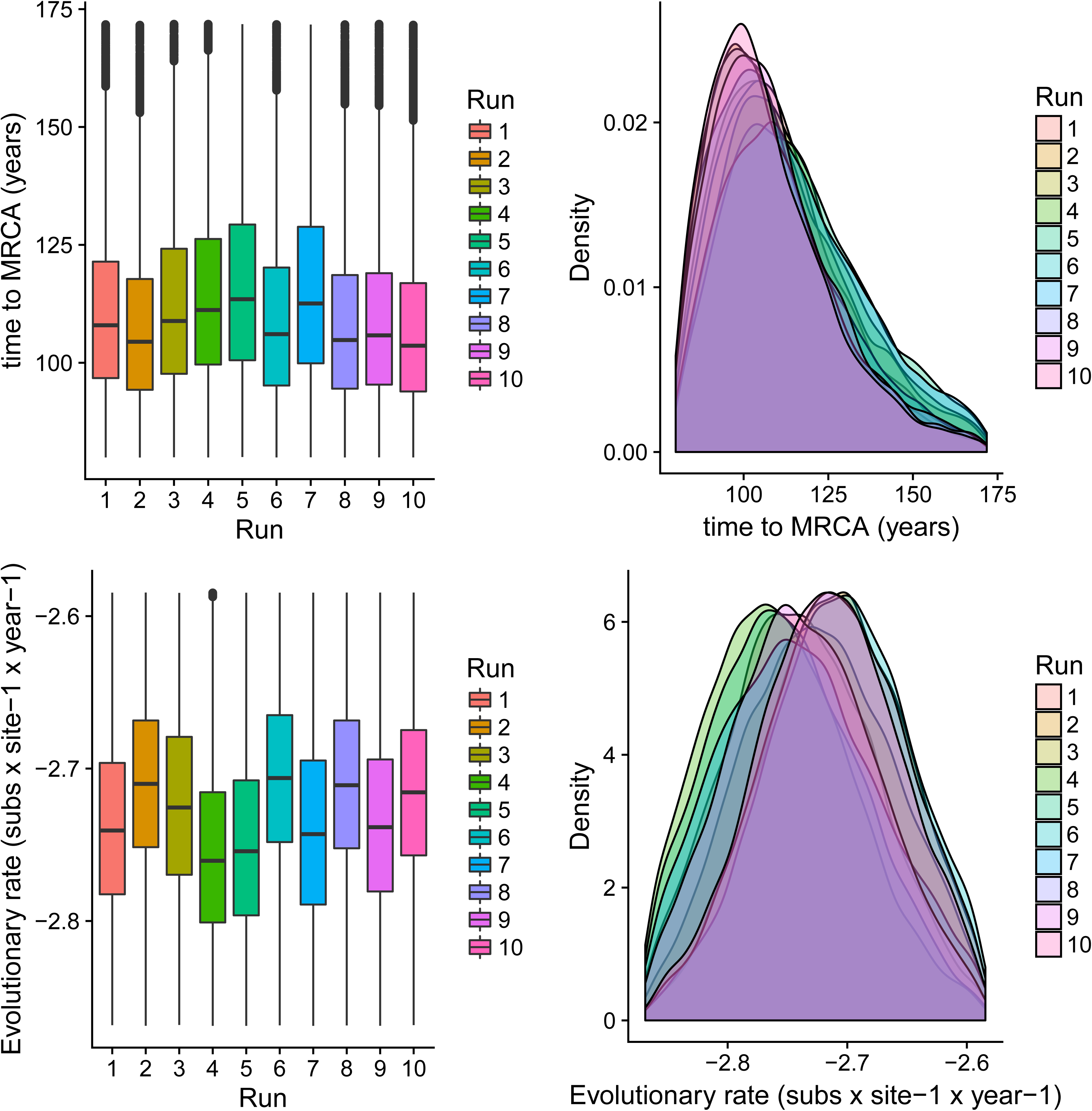
GI-16 genotype MRCA and evolutionary rate. Upper figure: Boxplot (left) and Densityplot of the MRCA posterior probability. Lower figure: Boxplot (left) and Densityplot of the mean evolutionary rate (expressed in base-10 logarithm) posterior probability. Results have been estimated performing ten independent runs based on sequences randomly sampled from the international database. The 95HPD intervals are reported for both figures.

The analysis of population dynamics revealed two main patterns: while the population size remained substantially stable between tMRCA and the 60s, a more fluctuating trend was observed thereafter (Figure 2). In the latter phase, two main peaks in the mean Ne·t value were consistently predicted around ∼1985 and ∼2005-2008. An additional minor rise was detected after 2010. Nevertheless, the variation in the population size slightly differed among runs and, more generally, the 95HPD intervals were generally broad over the whole considered period.

**Figure 2.**
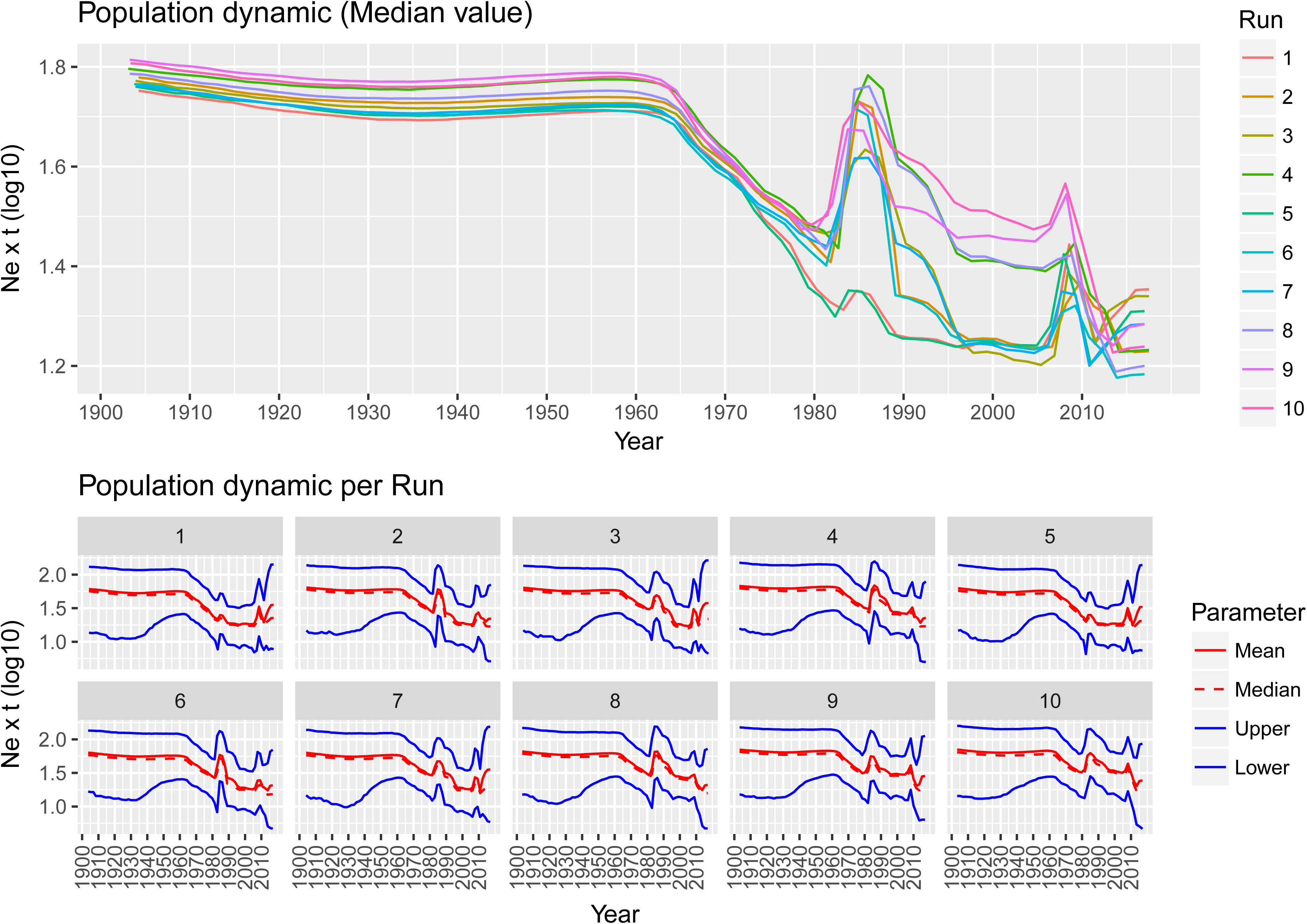
GI-16 genotype population dynamics. Upper figure: Mean relative genetic diversity (Ne·t) of the worldwide GI-16 population over time. The results of the ten independent runs have been color-coded. Lower Figure: Mean, median and upper and lower 95HPD values are reported for each run.

Phylogeographic analyses provided substantially constant results independently from the randomly generated database. Well supported migration rates were identified both among different continents, linking Asia with Europe, Middle East and South America, and within these regions, particularly in Asia and South America (Figure 3).

**Figure 3.**
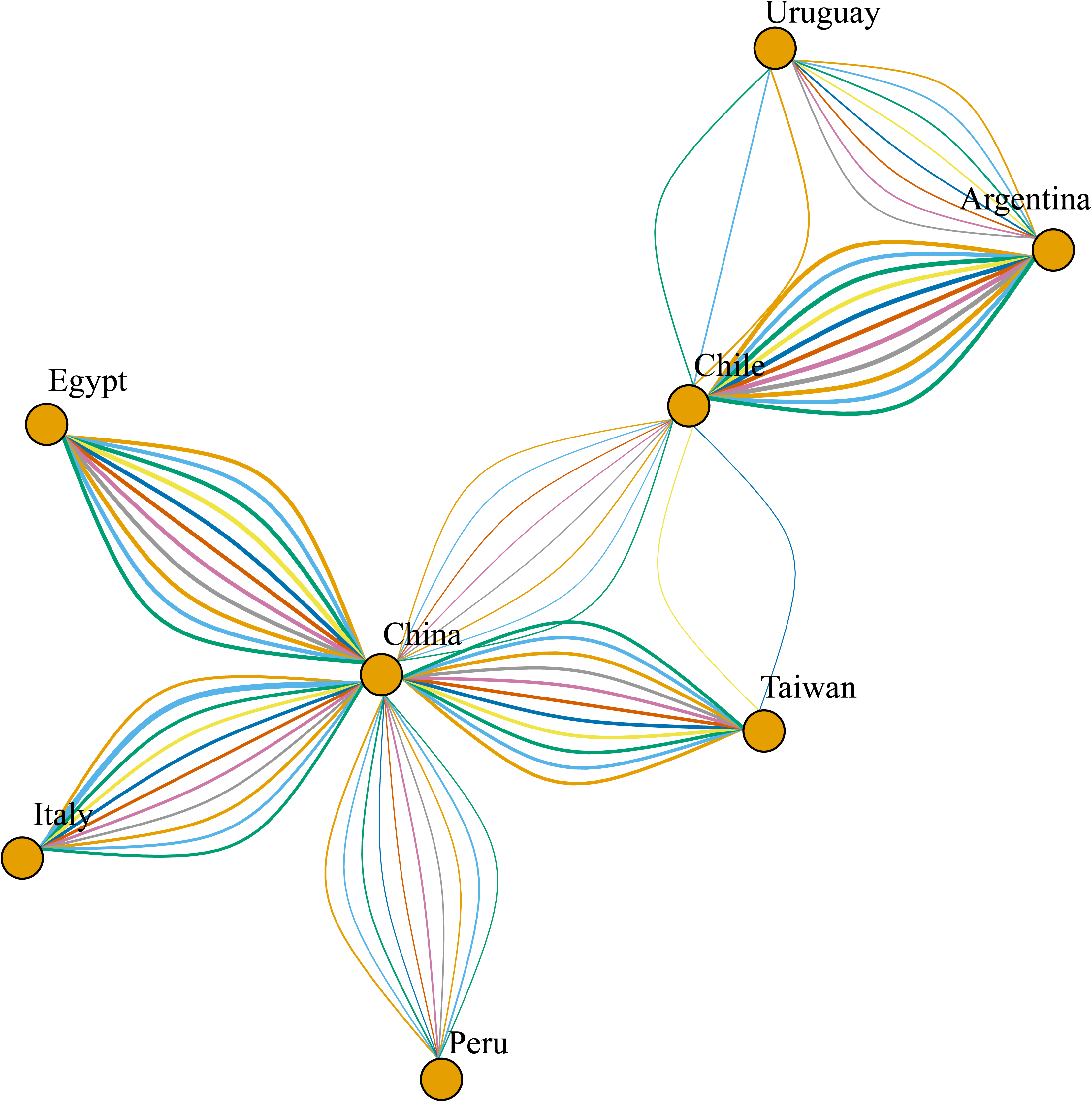
Network reporting the well supported migration routes(BF>3). GI-16 spreading path among different countries estimated using ten independent BEAST runs (color coded). The arrows size is proportional to the BF value.

The time-calibrated spreading process provided further information. Particularly, GI-16 lineage was limited to Italy until approximately the late 1980s, when a transmission to China was estimated (Figure 4). Thereafter, the virus migrated from China to other Asian countries (i.e. Taiwan) in the late 1990s, and in South America at the beginning of the new millennium, where it quickly spread among countries. An additional transmission from China to Middle East (Egypt) was predicted about in 2010. The location posterior probability was constantly high (i.e. >0.9) with the exception of just few ancestral nodes (S1 Fig), whose ancestral country could not be confidentially identified. For those particular ancestral nodes, differences were also observed among independent runs. However, this uncertainness is mainly on local base (e.g. China vs Taiwan or among South American countries) and does not affect the overall interpretation of GI-16 spreading pattern.

**Figure 4.**
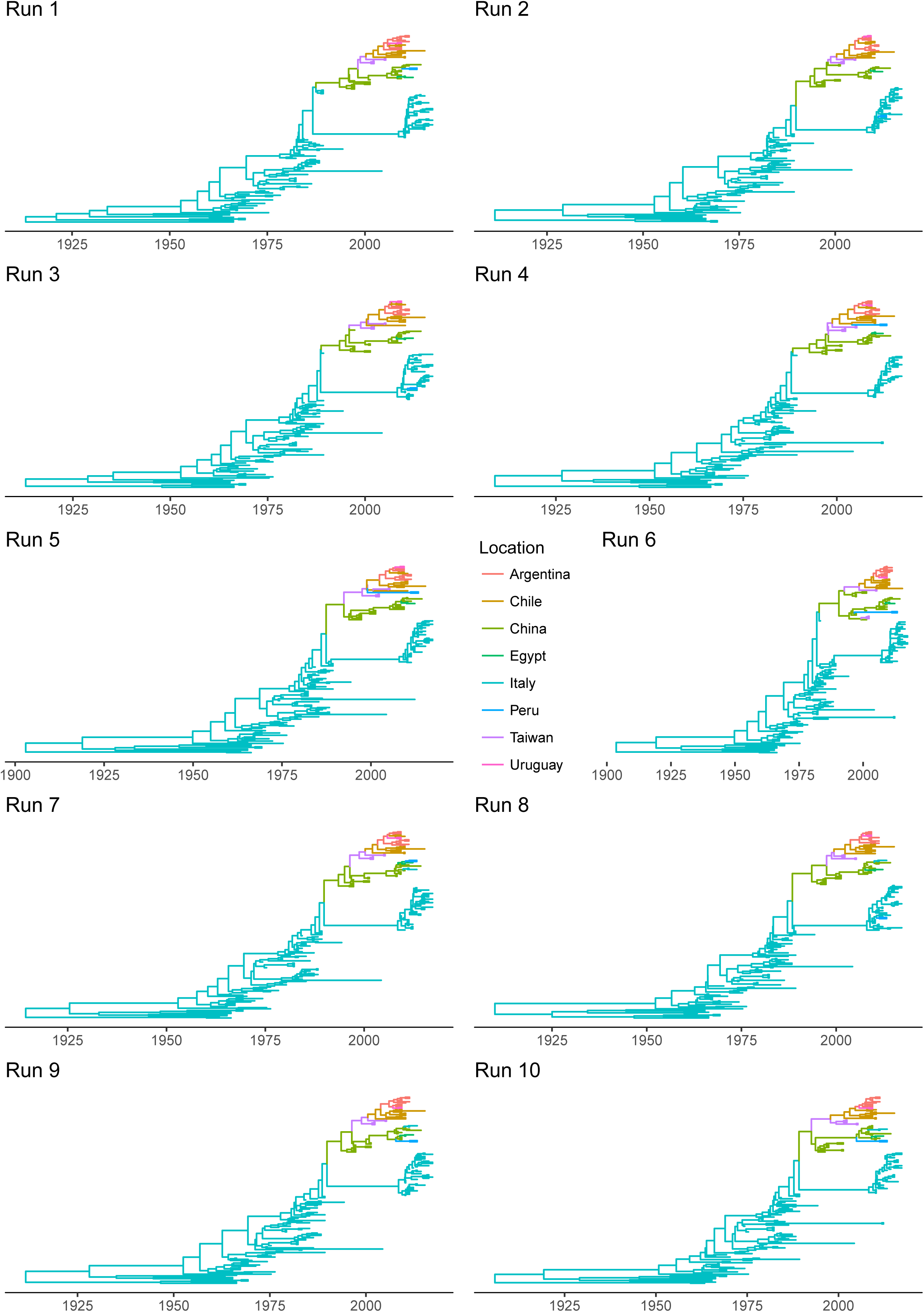
Time calibrated phylogentic trees. Time calibrated phylogentic tree obtained using ten independent BEAST runs are reported. The tree branches have been color-coded according to the location predicted with the highest posterior probability.

Almost all Italian strains collected after 2000 appear to descend from other strains detected in that country, supporting the persistent viral circulation. Two remarkable exceptions were represented by strain JQ290229.1|Infectious_bronchitis_virus|Italy|chicken|2011 and KP780179.1|Infectious_bronchitis_virus|Italy|chicken|2013, which were clearly related to Chinese ones and were therefore predicted to have been introduced from Asia (Figure 4). Because of the random sampling applied for the generation of the different databases, these two strains were not always included in all the performed analysis. Consequently, the relative viral migration from China to Italy has been reported only in a subset of the runs (i.e. runs 7 to 10, Figure 4).

### Phylodynamic analysis HVR12

The tMRCA and evolutionary rate were estimated to be 1964 (1943.84-1981.81 95HPD) and 3.16×10^-3^ (1.71×10^-3^-5.16×10^-3^). Population dynamic reconstruction evidenced a substantially stable population size, with an increase approximately in 2000 and after 2010 (S2 Fig).

Differently from HVR3, GI-16 origin was predicted in Slovenia. However, a high uncertainness featured this prediction, being Italy the potential root country. A migration from Europe to Asia was predicted in the ‘90s, potentially through independent introduction in China and Russia. Thereafter from China it spread to Taiwan, South Korea and South America (potentially to Argentina or Colombia) in the early 2000s. The following years were characterized by the viral spreading among South American countries. Independent introduction events from China to Peru, Egypt and Italy were observed around 2010 (S2 Fig). BF analysis evidenced the presence of several well supported migration rates among and within continents. Particularly, China (and Asia in general) appeared the most important nucleus of viral dispersion to other Asian and South American countries (S2 Fig).

## Discussion

The periodic emergence of new genotypes has been characterizing the epidemiology of IBV, continuously raising the bar for the implementation of effective control strategies. Although it has longly been accepted that the high evolutionary and recombination rates are the main drivers of IBV variability [3], the actual genesis of new genotypes remains elusive. In fact, these emerging variants are typically peculiar from a genetic perspective, which is incompatible with the recent differentiation from already known genotypes. Actually, the estimates of tMRCA of circulating lineages often largely backdate their first identification [23]. These evidences allow to speculate that many of the so called “emerging” genotypes could actually be quite ancient genetic groups that have not been correctly identified or classified yet.

The present study, reporting the evolutionary history of GI-16 lineage, supports this hypothesis. In fact, this lineage includes two genotypes that have been considered independent for a long time: the 624/I and Q1. The first one was firstly recognized in 1993 in Italy, albeit retrospective studies demonstrated the local circulation since 1963 [12], where it likely had a prominent role in IBV epidemiology. Nevertheless, its relevance appeared to gradually vanish during the 1990s. Almost simultaneously, a new genotype, often named Q1, emerged in China and was then reported in other Asian countries, Middle East, European and South American countries, being responsible of devastating economic losses [37]. The present phylodynamic analysis demonstrates a common origin of 624/I and Q1, which has been estimated approximately at the beginning or middle of the XX century, depending on the considered genomic region. Nevertheless, the lower sequence number and shorter time window covered by HVR12 database pose in favor of the HVR3 based analysis reliability. Anyhow, these results testify the long lasting undetected circulation of this genotype in Italy. The absence of adequate monitoring and diagnostic tools at that time clearly justifies this evidence, which could be realistically extended to most of current IBV genotypes. Unfortunately, the lack of sequences relative to the period before 1963 prevents the precise reconstruction of viral population dynamics in this time frame. However, the skyline profile, especially from the ‘60s onward, when actual data became available, shows that GI-16 was able to circulate and persist in Italy at low level until the ‘80s, when a remarkable increase in relative genetic diversity was observed. The reason behind this variation in the epidemiological scenario are hard to explain. However, a progressive intensification of poultry production, leading to higher animal density, turnover and presence of several concomitant stressing factors and co-infections could have created favorable conditions for the rise of acute respiratory infections like IBV. Similarly obscure are the causes of the GI-16 decline in the late 80s-early 90s. An upgrade in the vaccination strategies was applied in the ‘90s, through the introduction of 793B based vaccines [20]. A Mass plus 793B combined vaccination was proven effective in protecting against GI-16 induced disease [37], and could therefore justify the observed scenario. However, the new protocol was effectively introduced only on the last part of the decade. Consequently, even if it could have actually contributed to this genotype control, other factors, like changes in farm management or progressive increase of population immunity, must have played a role. Interestingly, an autogenous inactivated 624/I based vaccine (strain 794/83FO), administered at 1d of age, was introduced in the routine vaccination protocols of several farms. Although inactivated vaccines are traditionally considered less effective, a contribution in decreasing animal susceptibility and thus viral circulation can be suggested.

From the ‘90s onward this genotype followed a double fate. In fact, GI-16 was able to spread to China, which represented the main nucleus for a further diffusion. The introduction to South American countries (Figures 3 and 4) and the following spread among nearby countries are mirrored by a clear peak in viral population size (Figure 2) and correspond to the report of several and clinically relevant GI-16 induced outbreaks in these countries. The precise country of origin (China or Taiwan) and South American first-introduction country could not be established with confidence, being the results slightly different between S1 regions (i.e. HVT12 and 3) or among performed runs. Additionally, multiple introduction events involving different country pairs were observed (Figs 3 and 4). However, despite these variations due to the different sequences included in each dataset, the overall general spreading pattern appears clear and is characterized by a dominant eastward spreading from Europe to Asia and finally to South America.

Interestingly, although the migration rate was predicted to be low at the beginning of the GI-16 epizootiology (i.e. from Italy to China), the following steps were characterized by a much higher rate (Figure 5), likely reflecting the dense commercial connections among countries and the limited efficacy of implemented biosecurity measures [23]. The virus introduction from Europe to China could be ascribable to live animal movements, in particular of breeders that were imported to improve and/or substitute the local genetic lines. Additionally, it must be stressed that, even if the inferred link involved Italy and China, the GI-26 lineage was probably present before the ‘90s (undetected, improperly characterized or unsequenced) in several European countries, which could have played a relevant role in the eastward viral spread. However, extremely poor data reporting live poultry import and export from China are present and most of the countries considered in the present study declare no trade flows with China [14,18]. Therefore, other sources of viral spreading must be considered and the role of wild birds is often considered as a suggestive hypothesis. In fact, circulation of IBV related Gamma-coronaviruses in wild species, including migratory birds, has been reported, sometimes with high prevalence [38–40]. An involvement of these species in the spread of GI-16 can not be a priori excluded, especially over relatively short/medium distances (e.g within Europe and Asia or between these two regions) covered by recognized flyways. However, although the overlap between the East Asia/Australasia and East Asia/East Africa with the Pacific America and Central America flyways could explain the GI-16 introduction to South America, the extreme distance coupled with the absence of detection of this genotype in North America discount this hypotheses. Additionally, it must be stressed that GI-16 and other “wild type” genotypes have never been identified in wild birds and further studies will be necessary to properly and systematically investigate the role of wild birds in IBV epidemiology. In this background, other explanations like illegal or unreported animal trades appear more likely.

**Figure 5.**
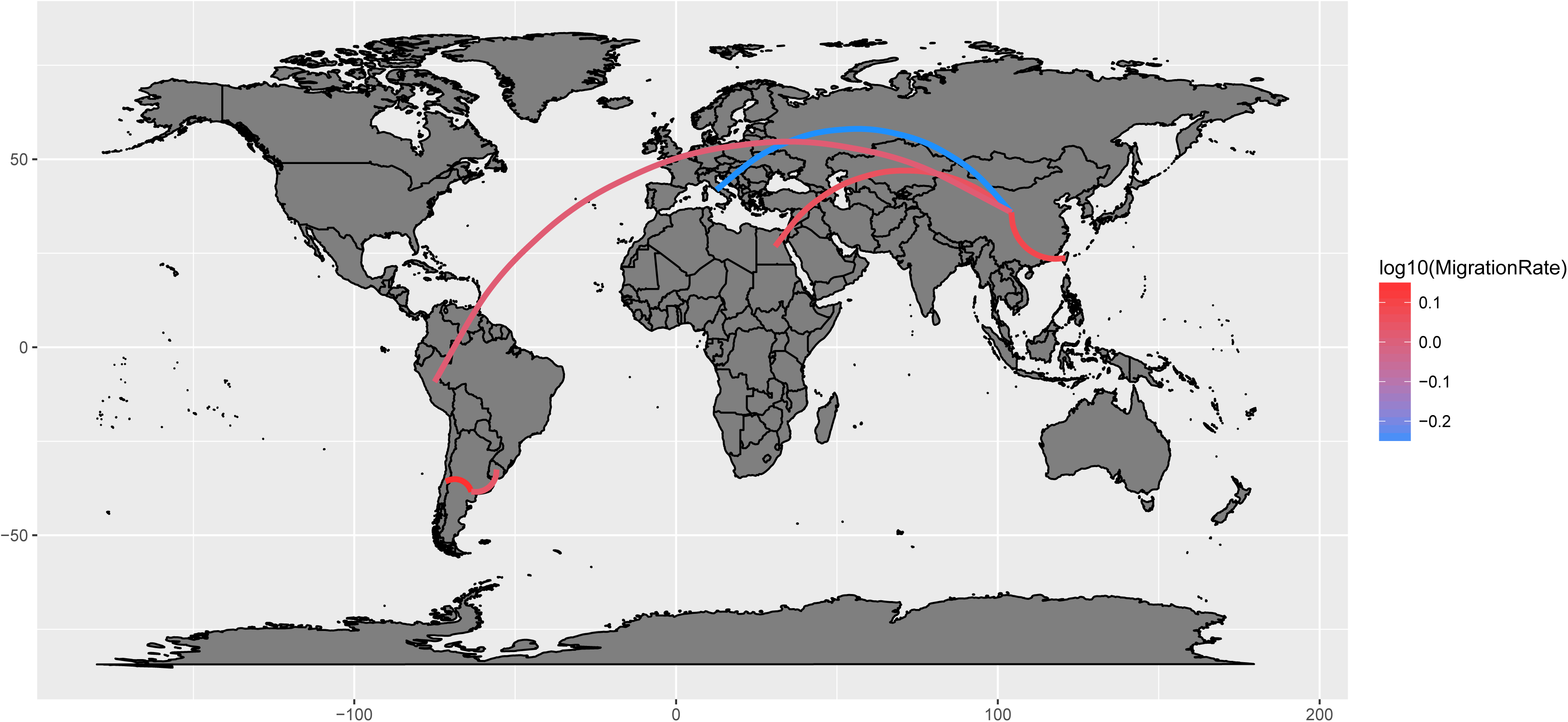
GI-16 genotype migration paths. Well supported migration paths (i.e. BF>3) among countries are depicted as edges whose color is proportional to the base-10 logarithm of the migration rate. The location of each country has been matched with its centroid.

Additionally, the limited number of available sequences could partially conceal more complex scenarios. For example, the presence of GI-16 in Slovenia [9] and Poland [10] in the ‘90s and in Russia [11] in the early 2000s could be suggestive of a more gradual migration from Mediterranean/Western Europe to Asia. Similarly, some intermediate steps in the Asian/American spread could have been missed for the same reason. Based on these results, the pivotal relevance of adequate monitoring activity and, when possible, retrospective studies appears clear for the understanding of viral disease epidemiology.

This concept is further supported by the depiction of the Italian scenario herein reported. Even though a limited number of Italian strains collected after 2010 were apparently introduced from China, the vast majority of recently sampled strains are direct descendant of the ancient “Italian 624/I” group. How this group was able to persist between the 1990s and the 2000s remains an unsolved question. A low-level circulation could clearly be advocated. Nevertheless, it must be stressed that the first identification of GI-16 was performed in the ‘90s. Additionally, three sequences of Italian GI-16, originating from research rather than diagnostic activity, were obtained in 1994, 1997 and 1999 [8,10,41] and similar strains were detected in the same period in a nearby country [9]. These evidences support a quite relevant but undetected genotype prevalence. Consequently, the effect of a remarkably lower sequencing activity in the 1990s compared to present days can not be under-emphasized and its consequences in terms of poor knowledge should be a reprimand to guide current monitoring activities planning.

Understandably, other factors could have played a role, like within-country viral evolution leading to the emergence of virulent strain, more fit for the Italian context. To investigate this issue, an ancestral state reconstruction of the amino-acid profile variation over time was performed using a Maximum-likelihood approach (S3 Fig). Albeit some amino-acid changes were detected in the clade including contemporary Italian strains, no distinctive mutations differentiating this group from their ancestors were identified, at least in the considered region. While the “local evolution” hypothesis appears lessened by this evidence, further studies will be necessary to investigate the potential role of other S1/genomic regions.

Finally, a variation in the competitive equilibrium between GI-16 and other pathogens or IBV genotypes (including vaccine strains) could have contributed to shape this genotype epidemiology. The introduction of GI-16 in South America represented an extraordinary epidemiological success from a viral perspective, as reflected by the increase in viral population size reported in the present study and by the outstanding clinical and economic impact in those countries [37]. On the other hand, the clinical impact of this lineage in Italy was negligible in the recent years. The competition with the highly prevalent QX genotype could have actually limited the potential of GI-16 in terms of prevalence and clinical relevance [20]. Additionally, the previously mentioned introduction of a combined heterologous vaccination in Italy (in South America flocks were vaccinated with Mass-based vaccines only) could have effectively protected the animals, avoiding the emergence of clinical signs and related economic losses.

Other studies have demonstrated the presence of relevant biological variations even within the same genotype [37]; the circulation of strains with peculiar pathogenic features could consequently be advocated. Remarkably, recent Italian and South American strains significantly differ in at least three protein positions (i.e. 47, 52 and 99 of the considered region). Interestingly, amino-acid in position 47 of South American strain was identical to the one featuring the ancient, and clinically relevant, Italian strains (S3 Fig) [42]. Even if suggestive, more detailed studied based on a reverse genetic approach will be useful to elucidate the actual influence of these residues on GI-16 virulence.

## Conclusions

Overall, the present study reconstructs the history and spreading dynamics of the GI-16 lineage from its origin to nowadays, demonstrating the complex network leading to its dispersal in Europe, Asia and South America. Above all, the ancient history of a previously considered “emerging” genotype is described and testifies the misconceptions that originate from the lack of a unified nomenclature and poor molecular epidemiology data generation and sharing. This shortcoming appears particularly relevant since the described scenario could likely be shared by many other IBV genotypes and pathogens in general.

## Supplementary material captions

**S1 Table List of sequences used in the present study and relative metadata.** List of HVR12 and HVR3 used in the study. Additional metadata including collection date and country are reported.

**S1 Fig. Ancestral location scatterplot.** Scatterplot representing the posterior probability of each ancestral location (color-coded) prediction over time. The results of ten independent BEAST run are reported.

**S2 Fig. HVR12 time calibrated phylogenetic.** Time calibrated phylogentic trees calculated using the HVR12. The tree branches have been color-coded according to the location predicted with the highest posterior probability (reported near the respective node). Left insert: Network reporting the well supported (BF>3) GI-16 spreading path among different countries (the arrows size is proportional to the BF value). Right insert: Mean and 95HPD relative genetic diversity (Ne x t) of the worldwide GI-16 population over time.

**S3 Fig. Acid ancestral state variation over time.** Maximum likelihood based reconstruction of amino acid ancestral state variation over time. Tips (sampled strain) amino-acids are reported as a color coded circle for each HVR3 position. The predicted amino-acids for each internal node (ancestral strains) are reported as a color coded pie-chart with slices proportional to the respective probabilities.

